# Detection and phylogeny of *Wolbachia* in field-collected *Aedes albopictus* and *Aedes aegypti* from Manila City, Philippines

**DOI:** 10.1101/2021.08.24.457456

**Authors:** Maria Angenica F. Regilme, Tatsuya Inukai, Kozo Watanabe

## Abstract

**Background:** *Wolbachia* is the most common bacterial endosymbiont of arthropods, such as the medically important *Aedes albopictus* and recent reports also detected in *Aedes aegypti*. Our study showed additional support for the presence of wolbachia in *Ae. albopictus* and *Ae. aegypti* using Wolbachia specific markers, *wsp*, and 16S.

**Main text:** This study collected 12 adult *Ae. albopictus* and 359 *Ae. aegypti* from 183 households in a dengue-prone area, Manila, Philippines, between June and September 2017. *Aedes* larvae (n = 509) were also collected from 17 water containers from 11 households. The DNA of the *Aedes* larvae and adults were screened for the presence of *Wolbachia* using the *wsp* and 16S markers, following optimized polymerase chain reaction (PCR) conditions, and sequenced. Our results showed that 3 out of 359 (0.84%) adult *Ae. aegypti* and 12 out of 12 (100%) adult *Ae. albopictus* were *Wolbachia* positive, whereas all larvae tested negative for *Wolbachia* (0/509; 0%). The *wsp* marker revealed six *Wolbachia-*positive *Ae. albopictus*, whereas the 16S marker showed *Wolbachia* in three *Ae. aegypti* and ten *Ae. albopictus*

**Conclusion:** Our results suggest that the utilization of two Wolbachia specific markers, *wsp* and *16S* demonstrated Wolbachia presence in individual *Ae. albopictus* and *Ae. aegypti*. The results of the Wolbachia infection in *Ae. albopictus* showed that the detected strains were from either supergroups A and B. Despite the low infection rate of Wolbachia in *Ae. aegypti,* our results demonstrated the presence of Wolbachia in field-collected *Ae. aegypti* supporting previous studies.

## Background

*Wolbachia* is a maternally inherited endosymbiotic bacteria, infecting 40% of arthropod species and some filarial nematodes [1-2]. Wolbachia is naturally present in medically important mosquito species, including *Aedes albopictus* [3-5] and *Aedes aegypti* [6-9] inhibiting the replication of arboviral pathogens [10]. Natural populations of *Ae. aegypti* were considered negative for *Wolbachia*, but recent studies have reported both positive [6-9;11-14] and negative [15-16] results for *Wolbachia* in field-collected *Ae. aegypti* among different countries. Due to these varying results, further validation of natural infection of *Wolbachia* in field-collected *Ae. aegypti* is needed.

The presence of natural *Wolbachia* strains in field-collected *Ae. aegypti* is important as it may affect future *Wolbachia* mass release programs, thereby influencing the successful invasion of *Wolbachia* strains from transfected mosquitoes into the natural population. Most studies have found that naturally occurring *Wolbachia* in *Ae. aegypti* are strains that are phylogenetically close to the wAlbB strain, which also infects *Ae. albopictus* [6-8;11]. *Ae. albopictus* is known to be infected with two *Wolbachia* strains, known as wAlbA and wAlbB, which belong to the supergroups A and B, respectively [17]. However, previous studies detecting *Wolbachia* in field-collected *Ae. aegypti* did not test *Ae. albopictus*, comparing natural Wolbachia strains present in *Ae. aegypti* with *Ae. albopictus* from the same localities at the same time, regarded as naturally infected is important to further characterize and validate natural Wolbachia infection in *Ae. aegypti*.

In this study, we detected *Wolbachia* in field-caught *Ae. albopictus* and *Ae. aegypti* using *Wolbachia*-specific DNA markers (*wsp* and 16S). We also identified and compared the *Wolbachia* strains and supergroups found in *Ae. albopictus* and *Ae. aegypti*. Our study provides further support in the presence of Wolbachia in field-collected mosquitoes.

## Main Text

Adults of 12 *Ae. albopictus* and 359 *Ae. Aegypti* were collected using commercially available mosquito light traps from 183 of 428 households randomly selected in Manila City, Philippines from June to September 2017. We found 17 water containers with *Aedes* larvae (*n* = 509) in 11 households. The adults (*n* = 371) and larvae (*n* = 509) were morphologically identified as *Aedes sp*. using a stereomicroscope using the keys of [18]. We extracted DNA using Qiagen AllPrep DNA/RNA micro kit and Qiagen DNA Blood and Tissue DNEasy Kits© (Qiagen, Hilden, Germany) in adult (*n* = 371) and larval (*n* = 509) samples, respectively.

*Wolbachia* was detected using: *wsp* (610 base pairs) with primer pairs *wsp* 81F (5′-TGGTCCAATAAGTGATGAAGAAAC-3′) and *wsp* 691R (5′-AAAAATTAAACGCTACTCCA-3′) [16] and 16S specific for *Wolbachia* (850 base pairs) with primer pairs *WolbF* (5′-GAAGATAATGACGGTACTCAC-3′) and *Wspecr* (5′-AGCTTC GAGTGAAACCAATTC-3′) [19].

For both *wsp* and 16S gene amplification, we followed the PCR protocol published in [7], in a final volume of 10 µl with 1 µl of the genomic DNA. We used the following components for the PCR reaction for both markers: 10x Ex Taq buffer, 25 mM MgCl2, 2.5 mM dNTP, 10 µM forward and reverse primers, 10% dimethyl sulfoxide, and 5 units/µl of Takara Ex Taq™. We included a positive control of a *Wolbachia-*positive *Culex sp*. and negative control of water in each PCR run. PCR products were analyzed in 1.5% agarose gel at 100V for 30 minutes. The criteria for a positive *Wolbachia* test were based on two successful amplifications per marker, *wsp*, and 16S.

We assembled and aligned the sequences for each marker using the CodonCode Aligner version 1.2.4 and MAFFT version 7. We checked all generated sequences for similarities with reference sequences from GenBank (NCBI, 2016) using Basic Local Alignment Search Tool–Nucleotide BLAST. All *wsp* and 16S, *s*equences were separately analyzed using DNAsp version 6.12.03 [20] to determine the number of haplotypes. We assessed the relationship of the *Wolbachia* strains of our study with representative sequences from different insect hosts by constructing a phylogenetic tree for the *wsp* and 16S sequences using PhyML 3.1 [21].

We screened all individual *Aedes sp*. larvae (*n* = 509) and adults (*n* = 371) for *Wolbachia* using the *wsp* and 16S. Larval samples showed no evidence of *Wolbachia* from either marker. However, *Wolbachia* was detected in 15 (4.04%) out of 371 *Aedes sp*. adult mosquitoes based on the *wsp* and 16S. A total of three (0.84%) were *Wolbachia* positive out of 359 *Ae. aegypti*, based on the *wsp* and 12 (100%) out of 12 *Ae. albopictus* (Figure 1), based on the *wsp* and 16S. The *wsp* detected *Wolbachia* in six *Ae. albopictus*, whereas the 16S detected it in three *Ae. aegypti* and ten *Ae. albopictus*. Among these *Wolbachia*-infected *Ae. albopictus*, 4 (26.67%) out of 15 were positive for both *wsp* and 16S. Confirmation of the species identification using the *cox1* revealed that the 12 adult mosquitoes were *Ae. albopictus*, whereas the remaining three were *Ae. aegypti* based on BLAST results with 100% identity match (Table 1).

**Figure 1.**
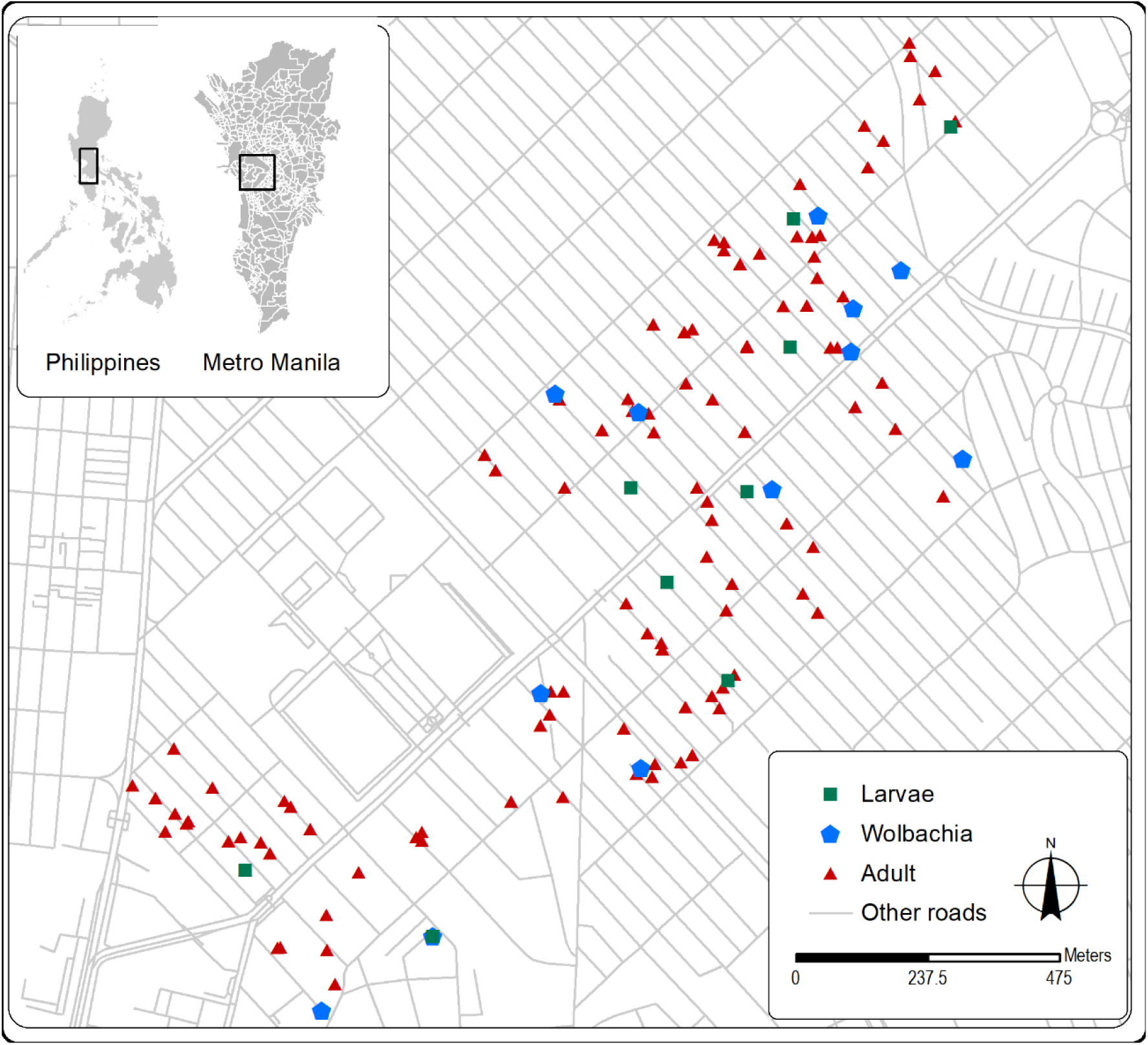
Spatial distribution of *Wolbachia*-positive households (n = 11) out of 428 households surveyed across a 2-kilometer road in highly urbanized area in Manila, Philippines.

**Table 1.** Summary of sampling data and Wolbachia detection results

|  | Total |
| --- | --- |
| Surveyed households | 428 |
| Households with adult Aedes mosquitoes | 183 |
| Households with larval Aedes | 11 |
| Collected Aedes mosquitoes | 371 |
| Aedes albopictus | 12 |
| Aedes aegypti | 359 |
| Collected Aedes larvae | 509 |
| Wolbachia positive |  |
| Larva | 0 |
| Adult Ae. albopictus | 12 |
| Adult Ae. aegypti | 3 |
| Households with Wolbachia-positive mosquitoes | 11 |

**Table 2.**
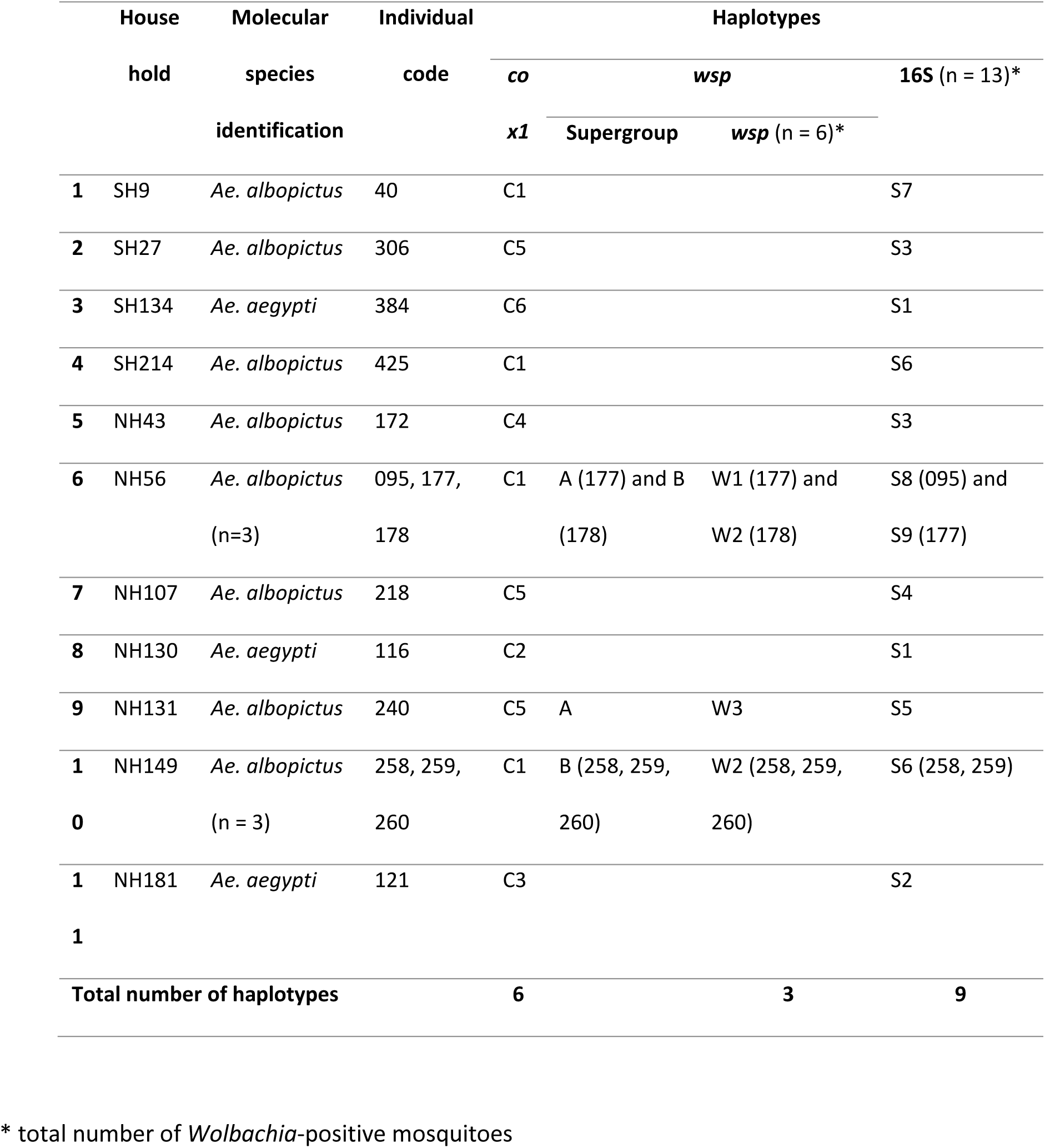
A detailed summary of the results of the Wolbachia infection in selected *Aedes* mosquitoes using *wsp* and 16S markers

The phylogenetic tree built using the *wsp* (Figure 2 A) revealed two major clades: supergroup A and supergroup B [15]. Two *Ae. albopictus* were infected with *Wolbachia* belonging to supergroup A and four with *Wolbachia* belonging to supergroup B. The haplotype sequences from supergroup A were grouped with the *Wolbachia* type strain A (wAlb A), identified in an *Ae. albopictus* host in the USA [17]. In contrast, the sequences from supergroup B clustered with the *Wolbachia* type strains from *Ae. aegypti* (*wAegB*) in the Philippines [6-7] and *Ae. albopictus* (*wAlbB*) in USA [7].

**Figure 2.**
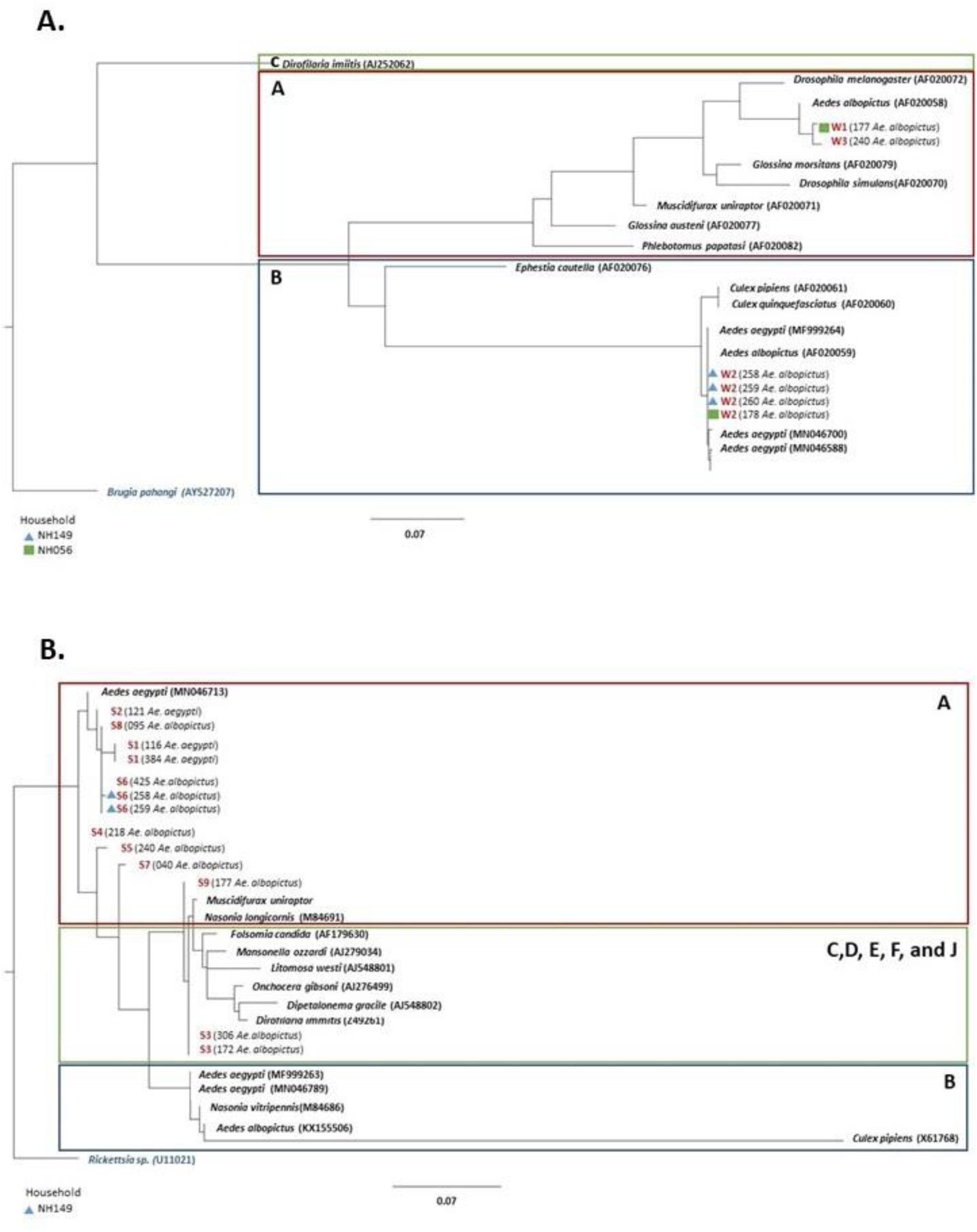
The maximum likelihood phylogenetic tree of *Wolbachia*-infected urban mosquitoes based on the **(A)** *WSP* sequences with reference sequences from supergroups A, B, and C and an outgroup of *Brugia pahangi* **(B)** 16S sequences with reference sequences from supergroups A, B, C, D, E, F, and J and an outgroup of *Rickettsia* sp.

In the phylogenetic tree constructed using 16S, 11 out of 13 sequences (84.62%) found in *Ae. albopictus* (n = 10) and *Ae. aegypti* (n = 3) were clustered within supergroup A (Figure 2 B). The remaining two (15.38%) found in *Ae. albopictus* were clustered together with the supergroups C, D, E, F, and J from insect hosts such as *Folsomia candida, Mansonella azzardi*, and *Dirofilaria immitis*.

We found that the *Ae. albopictus* were infected with both supergroups A and B of *Wolbachia* using the *wsp*, widely used for *Wolbachia* strain identification and phylogeny studies [17]. We found more *Wolbachia*-positive *Ae. albopictus* in supergroup B (4/6; 66.66%) than in supergroup A (2/6; 33.33%), as observed in a previous study [22] (China; 631/693; 91.05%). Our results support previous studies, and we observed two *Wolbachia* supergroups in *Ae. albopictus* collected from the same area at the same time.

We found a low prevalence of *Wolbachia* in adult *Ae. aegypti*, which is concordant with the previous results of [8] obtained from field-collected *Ae. aegypti* in Florida, USA and those of [7] conducted in Metro Manila, Philippines. In contrast, a high (>50%) prevalence rate was observed in the studies of [9] conducted in Malaysia and [8] conducted in New Mexico, USA. The low *Wolbachia* prevalence rate in *Ae. aegypti* may be due to the low density of the *Wolbachia* endosymbiont in *Ae. aegypti* [7]; this is supported by the previous metabarcoding studies of [8;11-12], which all reported a low number of sequence reads in the *Ae. aegypti* midgut, indicating a low density of *Wolbachia*. Although our results are limited since we did not quantitatively measure *Wolbachia* density, our 40-cycle PCR amplification method adapted from could infer that *Wolbachia*-positive *Ae. aegypti* might be present in Metro Manila, Philippines. Further *Wolbachia* detection studies are thus needed to affirm natural *Wolbachia* infection in field *Ae. aegypti*.

We acknowledge the uncertainties of *Wolbachia*-positive *Ae. aegypti* due to the 16S and the conventional PCR method, e.g., the false-positive rate. In light of this information, we were careful to ascertain positive *Wolbachia* in adult *Ae. aegypti*. Thus, we used the primers of [19], which are known to produce fewer false-positive results and negative detections of *Wolbachia*. We also performed repeated PCR tests of our *Wolbachia*-positive samples to ensure successful *Wolbachia* detection as defined in the previous study by [7]. In addition to conventional PCR, tests such as fluorescent *in situ* hybridization and whole genome sequencing can be used to provide strong support for the presence of *Wolbachia* in natural *Ae. aegypti* populations. Despite these limitations, our results provide additional evidence for *Wolbachia* detection in field-collected *Ae. aegypti*.

Our results were negative for *Wolbachia* in all the collected *Aedes* larvae (*n* = 509). One possible reason is that we found only 17 water containers during our field sampling. Since *Wolbachia* is maternally transmitted, all of its offsprings would also be negative for *Wolbachia* if the mother is negative for *Wolbachia*. More *Wolbachia* might be detectable in larval samples if we increase the number of water containers by setting up ovitraps. We also recommend detecting *Wolbachia* in representative larval samples from one water container. Future research may also explore the potential horizontal transmission of Wolbachia as observed in the study of [23].

## Conclusions

Our study showed the detection of Wolbachia in field-collected *Ae. aegypti* and *Ae. albopictus* in Manila City, Philippines. For *wsp*, we found six *Wolbachia* positive *Ae. albopictus* while the 16S showed three *Ae. aegypti* and ten *Ae*.*albopictus* positive for Wolbachia. We found 3 out of 359 (0.84%) in adult *Ae. aegypti* and 12 out of 12 (100%) adult *Ae. albopictus* while negative results were found in all larvae (05/509;0%). Our findings may provide additional support for the presence of *Wolbachia* in field-collected *Ae. aegypti* and its ability to invade and persist in the population.

## List of Abbreviations

PCR: polymerase chain reaction
DNA: deoxyribonucleic acid
*wsp*: Wolbachia specific primer

## Declarations

### Ethics approval and consent to participate

Not applicable

### Consent for publication

Not applicable

### Availability of data and materials

All data generated and/or analyzed during this study are included in this published article. All newly generated sequences are available in the GenBank database under the Accession Numbers. (Accession numbers have not yet been obtained at the time of submission, it will be provided during the review).

### Author’s contributions

MAFR and KW conceptualized and designed the experiment. MAFR and TI collected and identified the adult and larval samples for this study. MAFR and TI performed the *Wolbachia* detection and all the molecular experiments for this study. MAFR, TI and KW performed the data analysis. MAFR and KW wrote the manuscript. All authors read and approved the final manuscript.

## Acknowledgments

We are thankful to all the households that participated in our sampling collection of mosquitoes and larvae and to the village officials of Sampaloc, Manila, for their assistance and support. We appreciate the help of Johanna Beulah T. Sornillo of the Research Institute for Tropical Medicine, Philippines, for her technical assistance in the household sampling design. We want to thank Katherine Viacrusis for her service in the fieldwork and Mr. Micanaldo Francisco for constructing the sampling map. Our deepest thanks to Dr. Thaddeus M. Carvajal, Dr. Divina Amalin, Dr. Mary Jane Flores, Dr. Joeselle M. Serrana, and Jerica Reyes for their valuable suggestions and technical assistance. The authors would like to thank Enago (www.enago.jp) for the English language review. Maria Angenica F. Regilme is a recipient of the Japanese Government (Monbukagakusho) Scholarship from the Ministry of Education, Science, Sport, and Culture of Japan.

